# The turbulent soundscape of intertidal oyster reefs

**DOI:** 10.1101/2024.08.15.608049

**Authors:** Martin P. Volaric, Eli M. Stine, Matthew Burtner, Steven S. Andrews, Peter Berg, Matthew A. Reidenbach

## Abstract

Turbulence and sound are important cues for oyster reef larval recruitment. Numerous studies have found a relationship between turbulence intensity and swimming behaviors of marine larvae, while others have documented the importance of sounds in enhancing larval recruitment to oyster reefs. However, the relationship between turbulence and the reef soundscape is not well understood. In this study we made side-by-side acoustic Doppler velocimeter turbulence measurements and hydrophone soundscape recordings over 2 intertidal oyster reefs (1 natural and 1 restored) and 1 adjacent bare mudflat as a reference. Sound pressure levels (SPL) were similar across all three sites, although SPL > 2000 Hz was highest at the restored reef, likely due to its larger area that contained a greater number of sound-producing organisms. Flow noise (FN), defined as the mean of pressure fluctuations recorded by the hydrophone at *f* < 100 Hz, was significantly related to mean flow speed, turbulent kinetic energy, and turbulence dissipation rate (ε), agreeing with theoretical calculations for turbulence. Our results also show a similar relationship between ε and *FN* to what has been previously reported for ε vs. downward larval swimming velocity (*w_b_*), with both *FN* and *w_b_* demonstrating rapid growth at ε > 0.1 cm^2^ s^-3^. These results suggest that reef turbulence and sounds may attract oyster larvae in complementary and synergistic ways.

## 1 Introduction

Oyster reefs are dynamic ecosystems with a high degree of biotic and abiotic complexity. They provide habitat to a wide variety of organisms, which in addition to oysters includes snapping shrimp, xanthid crabs, polychaete worms, and a wide diversity of fish (Lenihan et al., 2001; Peterson et al., 2003; Rodney & Paynter, 2006). Oyster reefs also strongly impact local flow hydrodynamics (4), which can affect key ecosystem services such as reducing suspended sediment levels (5) and enhancing benthic metabolism (6,7). Due to these and other beneficial environmental impacts (8), there have been numerous efforts to restore oyster reefs, with successful restorations in both North America and Europe (9,10).

One metric that can potentially connect hydrodynamic and biological processes on oyster reefs is the soundscape, which arises from both physical and biological processes (11). It can be used as a proxy for the biodiversity and health of ecosystems (12,13), as well as to track the behavior of individual species (14). In recent years, researchers have investigated the soundscapes of a variety of marine systems, including coral reefs (12,15), rocky reefs (16,17) and oyster reefs (18,19). This work has focused primarily on temporal and spatial variations in the sound pressure level (SPL) across different frequency bands (e.g. Ricci et al. 2016; Salas et al. 2018; Monczak et al. 2019; Mueller et al. 2020). Sound frequencies above ∼150 Hz are often associated with marine biota, with frequencies 150 Hz < *f* < 1500 Hz indicative of fish chorusing (Ricci et al. 2016), and higher frequencies > 7000 Hz associated with snapping shrimp (Au and Banks 1998; Lyon et al. 2019). The lower frequency (*f* < 100 Hz) soundscape encompasses largely abiotic sounds, including breaking waves (24), rainfall (25), and anthropogenic sources such as ship noise (26). At these lower frequencies hydrophones also record pressure fluctuations resulting from current flow, turbulence, and wave orbitals. Although not a true sound wave, this so-called “flow-noise” (FN) contains a wealth of information about the hydrodynamic environment (Wenz 1962; Strasberg 1979; Bassett et al. 2014; Auvinen et al. 2019).

Flow noise and sound waves are similar in that they create periodic pressure variations over time; FN typically occurs at lower frequencies, but that is not a distinguishing characteristic. Instead, the critical difference is that FN pressure variations move with the water flow, which is generally on the order of centimeters per second, whereas sound wave pressure variations move as compression waves that propagate at the speed of sound in water, which is about 1500 m s^-1^. FN is a near field effect, meaning that it is detected less than one wavelength from where it is produced, while true sound waves are in the far field, meaning that they are detected several wavelengths from where they are produced. Near field effects share the behaviors that their absorption at one location feeds back to the objects that produced them at a different location, their phase relationships often differ from far-field waves (e.g. displacements in pressure and velocity are in-phase for far-field sound waves but often out-of-phase for near-field effects), and their strengths die off very rapidly with increasing distance (31).

### 1.1 Oyster larvae behavioral responses to sound and turbulence

When transitioning from pediveliger larvae to spat, Eastern oyster (*Crassostrea virginica*) larvae first make their way to a suitable settlement location (Poirier et al., 2019) by descending to the seafloor, moving horizontally in search of potential settlement substrates, and then either affixing to a substratum or ascending to rejoin the water column (Hadfield & Paul, 2001; Meyer-Kaiser et al., 2019). *C. virginica* larvae likely expend energy at a high rate while propelling themselves, given that the reward of a good site (i.e., heightened chance of settlement or avoidance of predators) offsets the risk and expenditure of the search (Fuchs et al., 2015b). The larvae use an assortment of chemical, physical, and auditory cues from the surrounding environment to make these decisions (Meyer-Kaiser et al., 2019). Physical cues are sensed by *C. virginica* larvae using a structure known as the statocyst, which detects gravity, turbulence, vorticity, acceleration, and orientation (Fuchs et al., 2013; Wheeler et al., 2015). Turbulence within the water column stimulates active downward propulsion of *C. virginica* larvae, increasing the likelihood of settlement (Fuchs et al., 2013). Statocysts detect sound for different mollusks such as cephalopods (Kaifu et al., 2008), and have been hypothesized to be the primary mechanism for sound detection in adult oysters (Charifi et al., 2017). Because the statocyst can sense fluid particle motion and therefore particle accelerations caused by sound waves, it is also believed to help larvae detect elements of the aquatic soundscape (Lillis et al., 2014b), allowing larvae to orient themselves according to auditory cues when navigating towards a potential settlement site (Lillis et al., 2013b).

Additional evidence for the soundscape influence on larval settling arises from research using manipulated sounds. Using calibrated hydrophones, researchers recorded the sounds of Australian flat oyster (*Ostrea angasi*) reefs and broadcast them at three sites of varying background noise levels using underwater speakers set to parameters representative of the natural habitat (Williams et al., 2022b). Sites in which the reef soundscape was audibly enriched (and therefore the reef soundscape was elevated above the background soundscape) had significantly higher *O. angasi* larval settlement than those with medium and high background noise. For *C. virginica*, larvae have been shown to preferentially swim towards recordings of reefs vs. unstructured bottoms (Lillis et al., 2013), possibly due to differences in SPL at *f* > 2000 Hz (37).

### 1.2 Study objectives

As previous studies have shown that both turbulence and sound promote larval settlement but have only investigated them separately, a better understanding of the linkage between hydrophone soundscape recordings and turbulence is needed. In this study, we paired marine soundscape recordings with concurrent acoustic Doppler velocimeter (ADV) turbulence measurements at three intertidal sites: a natural oyster reef, a restored oyster reef, and a bare mudflat. Our objective was to establish the link between oyster reef soundscapes and hydrodynamics, specifically the role that turbulence plays in generating near field pressure waves that may be used as physical cues by oyster larvae when initiating settlement behaviors.

## 2 Data and Methods

### 2.1 Study site

The study site consisted of two intertidal *Crassostrea virginica* oyster reefs – a mature, natural reef and a restored reef constructed by The Nature Conservancy (TNC) – and a nearby intertidal bare mudflat located within the TNC monitored Hillcrest Oyster Sanctuary along Virginia’s Eastern Shore (Fig. 1). This sanctuary is within the Virginia Coast Reserve, part of the National Science Foundation’s Long-Term Ecological Research network. The natural reef measured 240 m^2^ in size (22.5 m long x 10.5 m wide) and had a mean oyster density of 103 ± 27 oysters m^-2^ (mean ± SE, n = 12). The restored reef was established by TNC in 2010 (38) using piled oyster shell and measured 2275 m^2^ in size (35 m long x 65 m wide). It had a slightly greater but not significantly different oyster density than the natural reef, with a mean density of 128 ± 26 oysters m^-2^ (n = 20). All density counts were for individuals > 50 mm in length using 0.5 m x 0.5 m quadrats placed randomly on the reef. The elevations of the three sites ranged from −0.35 to 0.07 m relative to local mean sea level (39), and the depth at high tide was approximately 1-1.5 m. The observational setup is depicted in Fig. 1A (shown for restored reef).

**Figure 1.**
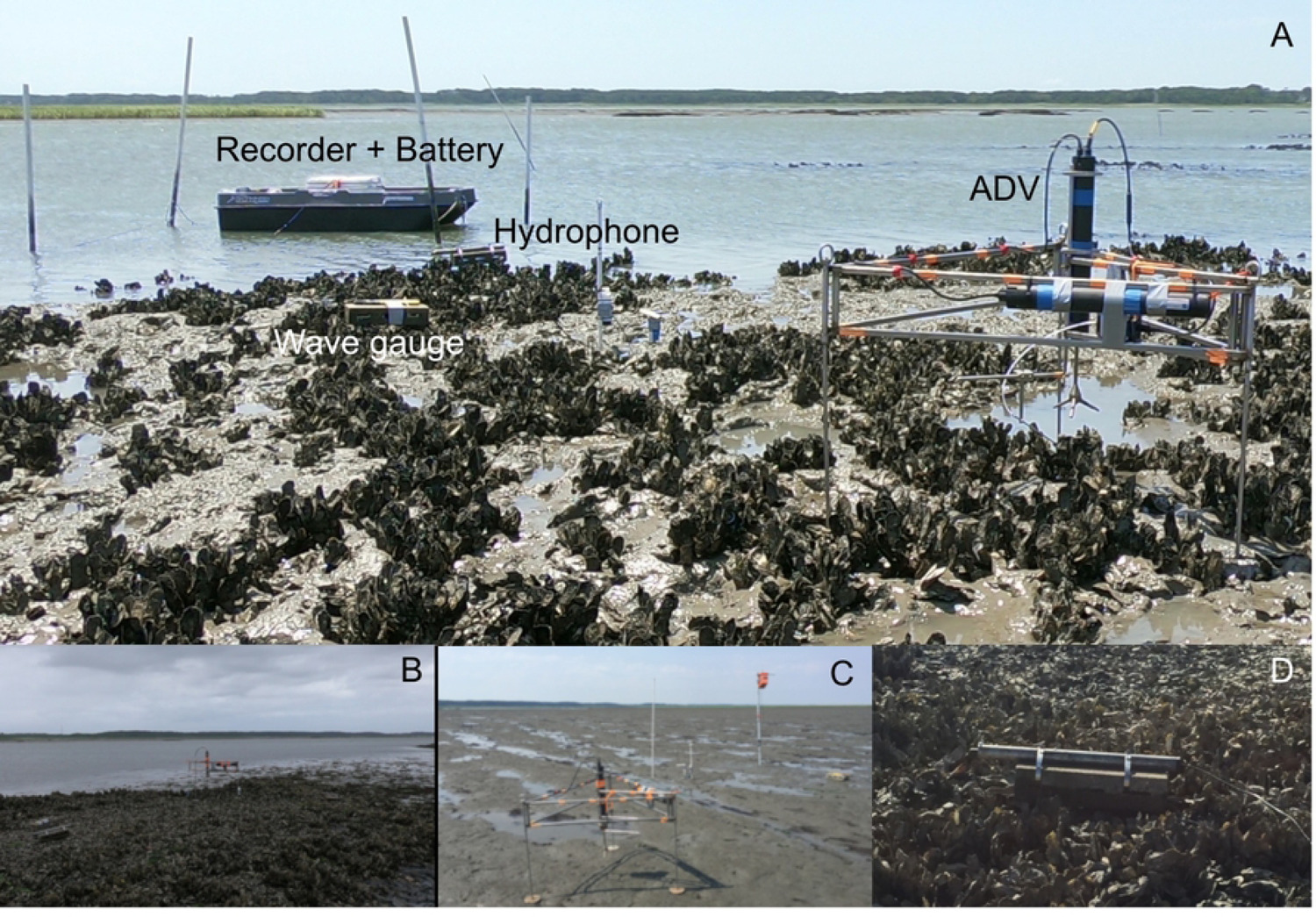
(A) Restored reef depicting observational setup which include a hydrophone attached to recorder and battery, wave gauge, and acoustic Doppler velocimeter (ADV). (B) A natural reef and (C) mudflat were also sampled with the same instrument array. (D) Close up showing hydrophone extending out of metal tubing a fixed distance above the reef. These images depict these intertidal sites during emergent conditions, but data was only used when they were submerged.

### 2.2 Sound recordings

Sound recordings were made using an omnidirectional Aquarian Audio H2A hydrophone (sensitivity: −180dB re: 1V/µPa, linear range 20Hz - 4KHz) connected to a Zoom H6 portable recorder. The hydrophone was secured to the end of a metal tube strapped to a cinderblock placed approximately 10 cm above the reef (Fig. 1D). The sound recorder was powered by an external DC battery that allowed for ∼48 hours of continuous recording, made at either 16 bit/96 kHz (oyster reefs) or 16 bit/44.1 kHz (mudflat), before being later downsampled to 25kHz. The portions of recordings during which the hydrophones were exposed to air during mid to low tide were discarded, resulting in recordings of 20 high tide periods (6 from the natural reef, 8 from the restored reef, and 6 from the mudflat), encompassing more than 200 hours of sound recordings from the sites.

Pressure fluctuations recorded by the hydrophone were converted into dB through laboratory calibration of known 80 dB and 100 dB sounds using a reference sound pressure of 1 μPa, and then split into three frequency (*f*) bands using high and low pass filtering. The lowest frequency band (FN) consisted of data at *f* < 100 Hz, the medium frequency band (SPL_MID_) at 150 Hz < *f* < 1500 Hz, and the high frequency band (SPL_HI_) at *f* > 2000 Hz. Mean FN, SPL_MID_ and SPL_HI_ were then calculated over 15 min intervals.

The dB scale represents a log-transformed value of the amplitude of pressure fluctuations recorded by the hydrophone. For certain comparisons with hydrodynamic results, we converted FN from dB to Pa with:

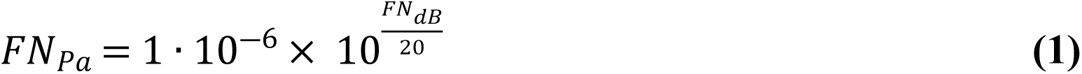

### 2.3 Turbulence measurements and data analysis

Turbulence measurements were made using a Nortek AS Vector^©^ acoustic Doppler velocimeter (ADV). This instrument makes 32 Hz recordings of the three velocity components (*u*, *v*, *w*) within a cylindrical measuring volume, 14 mm in diameter and 14 mm in height, with the volume set to 10 cm above the reef defined as the tops of oyster shells. The 32 Hz data from the ADV were averaged to 8 Hz, followed by a 2-step coordinate rotation in which the mean velocity was assigned to the horizontal component *u*, and the mean transverse velocity *v̅* and mean vertical velocity *w̅* were set to 0. Coordinate rotations and mean velocities were computed for each 15 min time interval.

Turbulent kinetic energy (*TKE*) was calculated as:

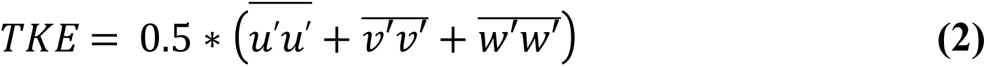

where *u’*, *v’*, and *w’* are the turbulent fluctuations about the rotated mean velocities and the overbars indicate time averaging over 15 min intervals.

The turbulence dissipation rate (ε) over each 15 min interval was calculated from a one-dimensional spectral equation of *w’* (Shaw et al., 2001; Reidenbach et al., 2006):

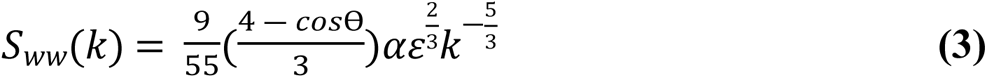

where *S_ww_*(*k*) is the power spectral density as a function of wave number *k*, Ɵ is the angle from the direction of mean flow and is equal to 90°, and α = 1.56 is the Kolmogorov constant for velocity. Further explanation of ε calculation is provided within Reidenbach et al. (2006).

The turbulence dissipation rate can be used to estimate the bed shear stress (*τ_b_*), assuming a balance of production and dissipation of *TKE* within the constant stress region of the boundary layer (42). The ‘law of the wall’ can be used to relate the friction velocity, *u_*_*, to the *TKE* dissipation rate as *u*_∗_ = (*ɛκz*)^1/3^, where *κ* =0.41 is von Karman’s constant, and *z* is the elevation above the seafloor. *τ_b_* was then calculated as:

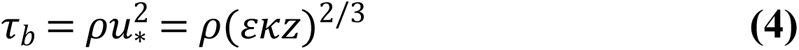

where *ρ* is the density of seawater at 25 °C and a salinity of 32 (1022 kg m^-3^).

### 2.4 Wave measurements

Pressure at the seafloor was recorded at 8 Hz using an RBR^©^ wave gauge. Significant wave height H_s_, defined as the mean of the highest one-third of waves, was calculated as:

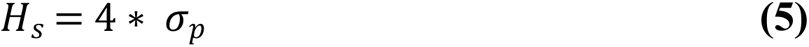

where *σ_p_*is the root mean squared error of a linear fit, used to remove tidal effects, of the pressure signal over each 15 min interval (43).

## 3 Theory

We also investigated the link between hydrodynamics and the underwater soundscape using hydrodynamic theory. To do so, we assumed that tidal flow over a reef can be treated as the sum of the mean flow and well-developed isotropic turbulence that is characterized by a continuous spectrum of eddies with energy flow from larger to smaller ones.

In the water column, the largest eddies are on the size scale of the water depth, *d*. However, velocities were measured at an elevation z = 0.1 m above the reef, therefore the largest bed-generated eddies that we typically observed with our ADV turbulence measurements were constrained by this 0.1 m elevation. The length scale of unstratified wall-bounded shear flows is proportional to the distance from the boundary, *L* = *κz*, where *κ* = 0.41 is von Karman’s constant (44,45), giving the scale of our largest observable eddies as *L* = 0.041 m. At the other extreme, the smallest eddies are on the scale of the Kolmogorov length (*η*):

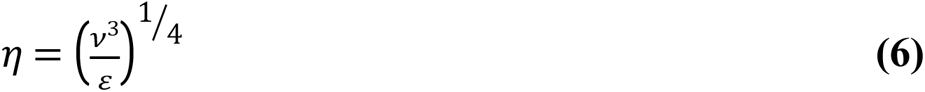

where *ν* is the kinematic viscosity, which is about 10^-6^ m^2^/s for water. The Kolmogorov length is approximately equal to the length at which the Reynolds number is equal to 1 and varied from ∼0.2 to 2 mm at our sites depending on flow rate.

Between these length scales, called the inertial subrange, the turbulent kinetic energy is partitioned amongst the differently sized eddies according to the Kolmogorov energy spectrum:

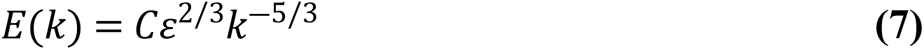

where *C* =1.56 is the empirical Kolmogorov constant for velocity and *k* is the wavenumber measured in radians per meter, with *k* = 2π/*L* where *L* is the eddy size. Over the inertial subrange, *k* varies between *k*_*min*_and *k*_*max*_, with:

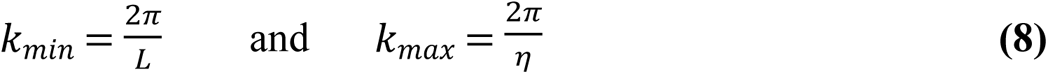

The spectrum of pressure fluctuations can be computed for isotropic turbulence over the inertial subrange (46):

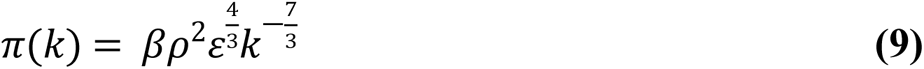

where *π*(*k*) is the pressure as the function of wavenumber and *β* = 2.97. Integrating this spectrum over all wavenumbers with significant turbulent energy leads to the mean square pressure fluctuation:

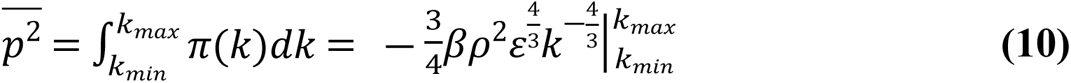

The result is approximated using the fact that *L* >> *η*, which makes the *η* contribution negligible. The root mean squared pressure fluctuation caused by turbulence can then found by substituting *k_min_*into Eq. 7 and taking the square root:

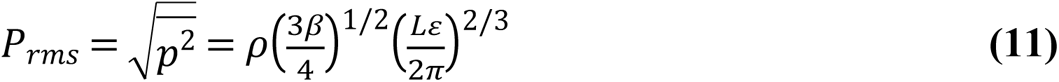

These pressure fluctuations propagate with the water flow, including both mean flow and turbulent flow, which then produces low frequency pressure variations that are recorded by the hydrophone. It is these fluctuations that we define as FN.

## 4 Results

### 4.1 Soundscape

Fig. 2A presents an example sound spectrogram from one high tide period at the restored reef. This distinctive canoe-shaped spectrogram is typical of what we observed at all three sites. There was little sound energy in the 1000 - 3000 Hz range at the beginning and end of each high tide period, when the water was shallow, and greater sound energy in this range at peak high tide when the water was deeper. Fig. 2B shows more detail by presenting sound energy spectra for a mid-tide time (red) and the high-tide time (black), again revealing a frequency band with low sound energy. In both panels, the sounds at the lowest frequencies (f < 100 Hz) represent FN, while those at higher frequencies represent true sounds (SPL). The higher frequency sounds, from 1 kHz upward, were predominantly produced by snapping shrimp. This is supported by the fact that these sounds were not continuous but were separate distinct “clicks”, which can be seen in the spectrogram shown in Fig. 2C. The intensity also decreases with frequency as approximately *f*^-6.3^ (blue line in Fig. 2B), which is a much steeper decrease than would be expected for sound that is produced from turbulence (either *f*^-1.3^ or *f*^-3.5^, from Rubinstein and Zhou, 2000). Similar sounds were recorded at the mudflat and both oyster reefs, all of which support snapping shrimp populations.

**Figure 2.**
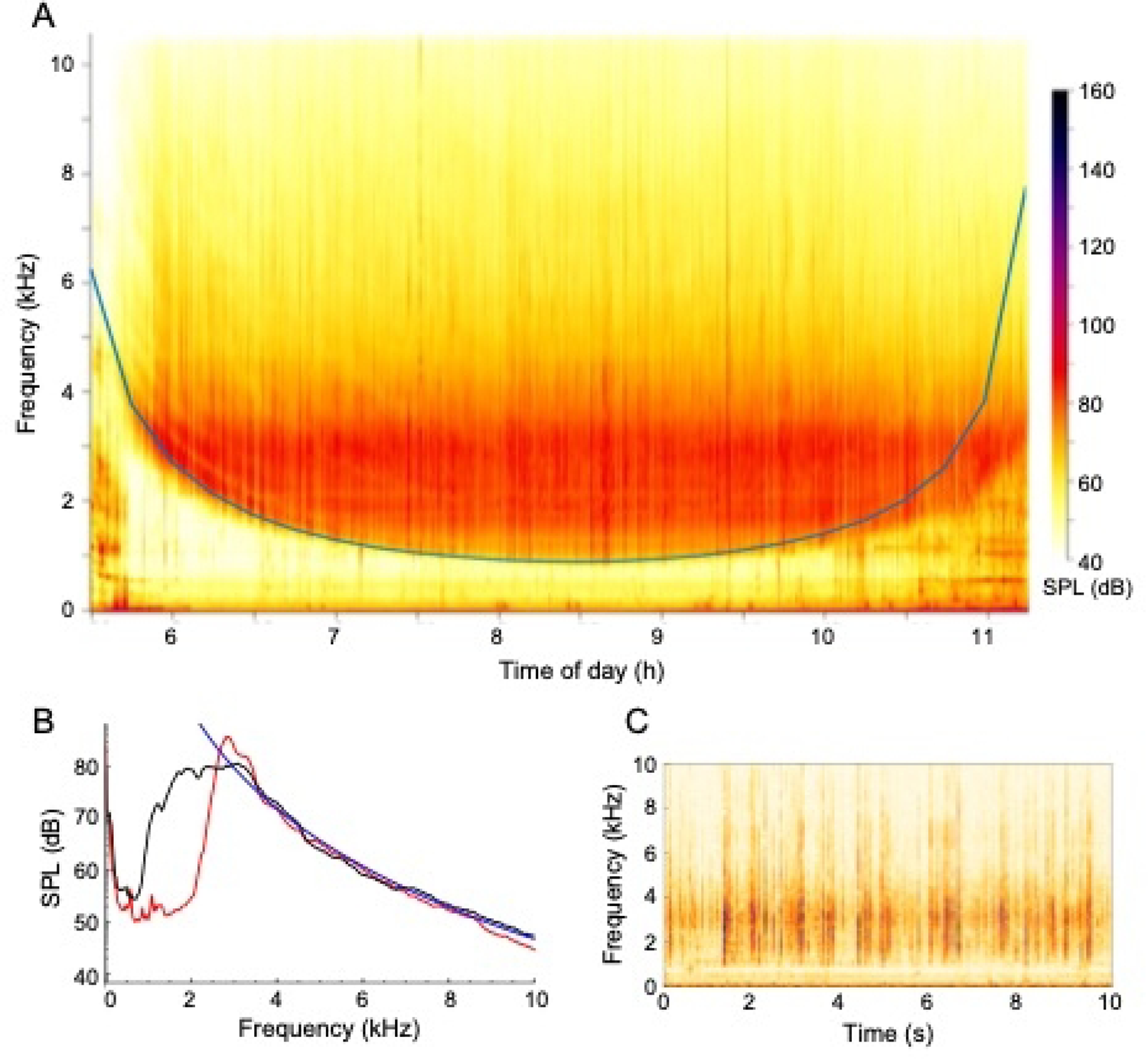
**(A)** Example power spectra over 1 high tide period from restored reef taken the morning of June 14, 2018. This spectrogram is representative of all three sites, with SPL at higher frequencies showing a consistent relationship with depth, resulting in distinctive “canoe-shaped” spectrograms in which there was very little sound energy in the 1000 - 3000 Hz range at the beginning and end of each tidal period. (B) Sound power spectra from the same data set, shown at times 6 h in red, and 8.5 h in black. The blue line is a fit in which SPL, in Pa, is proportional to *f*^-6.3^. (C) A 10 s long spectrogram from the same data set from time 8.5 h, showing that the sound arises from a series of distinct clicks, not from a continuous noise source.

The distinctive curve in Fig. 2A arose from water depth effects. More precisely, shallow water acts as an acoustic waveguide that channels sounds between the sediment and water surface boundaries (47). Such waveguides have a cut-off frequency (i.e. a minimum frequency that can be transmitted) which is equal to:

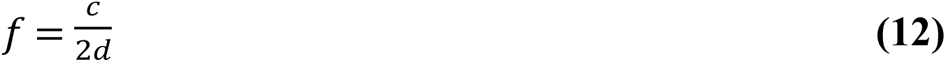

Fig. 2A shows this cut-off frequency, which agrees very closely with the experimental data. The primary differences occur during times with increased wave activity, such as during hours 10 to 11.5 in Fig. 2A, which disturb the upper boundary and decrease the waveguide effect.

Fig. 3 compares 15 min mean values for FN, SPL_MID_, and SPL_HI_ for the three sites. It shows that they are qualitatively similar, although one-way ANOVA tests for all three sound bands show statistically significant (F > 36, p < 0.001) differences between the 3 sites for all 3 sound bands. In particular, the restored reef exhibited higher SPL_MID_ and SPL_HI_, which we suspect arose from its substantially larger area.

**Figure 3.**
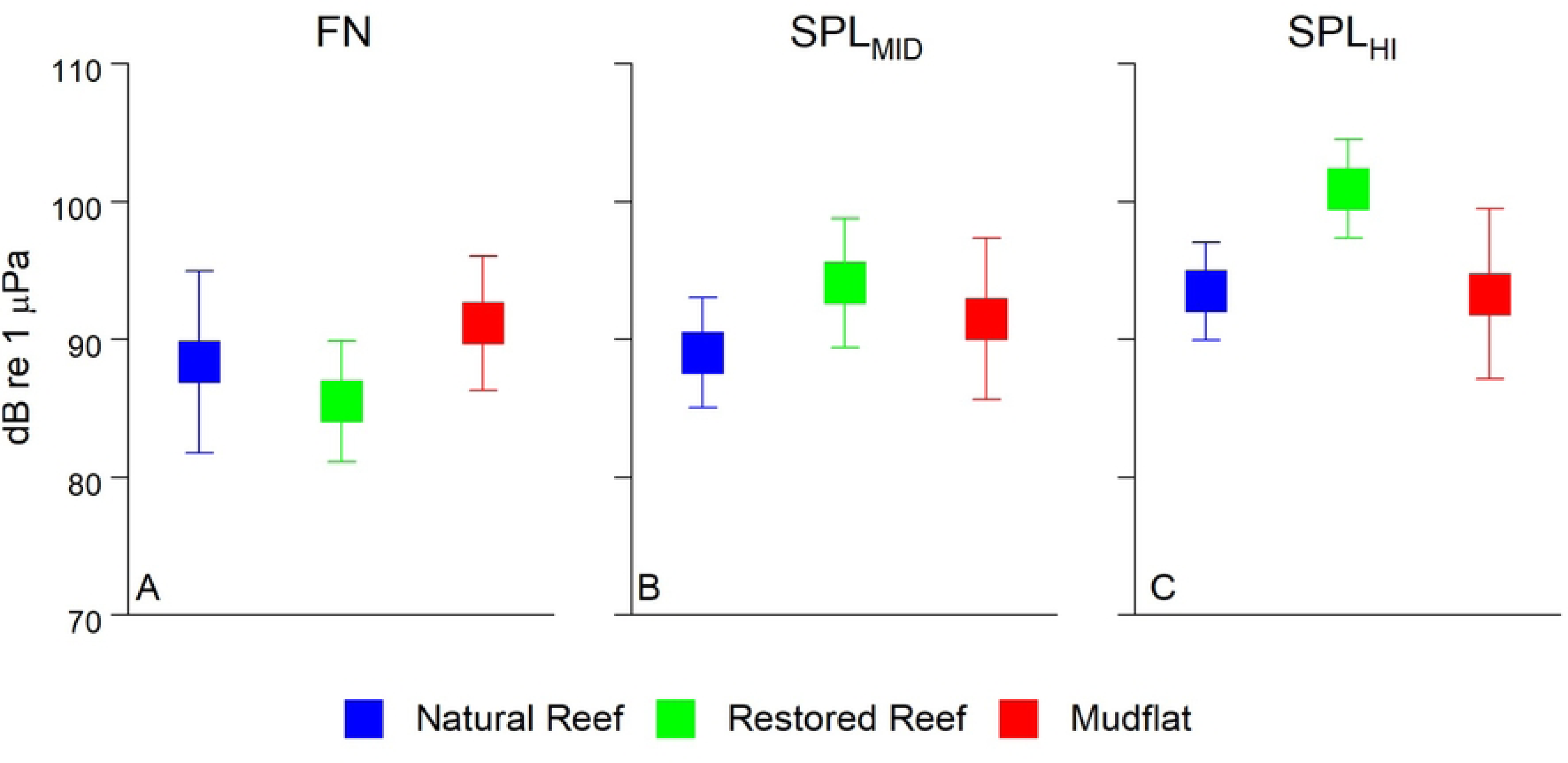
Mean FN or SPL for each of the three frequency bands for each site. Datapoints represent mean of 15 min averages ± SD. (A) FN < 100 Hz, (B) 150 Hz < SPL_MID_ < 1500 Hz, (C) SPL_HI_ > 7000 Hz.

### 4.2 Turbulence and hydrodynamics

Fig. 4 depicts a portion of a typical data set, in this case collected at the mudflat and including a thunderstorm that occurred between times 20.25 and 20.75 h. Fig. 4A shows raw ADV data, before coordinate rotation, where it is seen that the rising tide produced negative flow along *u*, positive flow along *v*, and negligible flow along *w*. Dots show the total flow speed, *U*, calculated as the root-mean-squared velocity magnitude. The flow direction reversed during the storm, presumably due to wind effects, and then returned afterward. Fig 4B shows that the depth increased with the rising tide, and that significant wave height was minimal before and after the storm but up to about 20 cm during the storm. Fig 4C shows the impact of the flow and storm on FN. It exhibited a spike during the storm and was also greater at the beginning of this window, when flow velocities were higher, and lower at the end, when velocities were lower.

**Figure 4.**
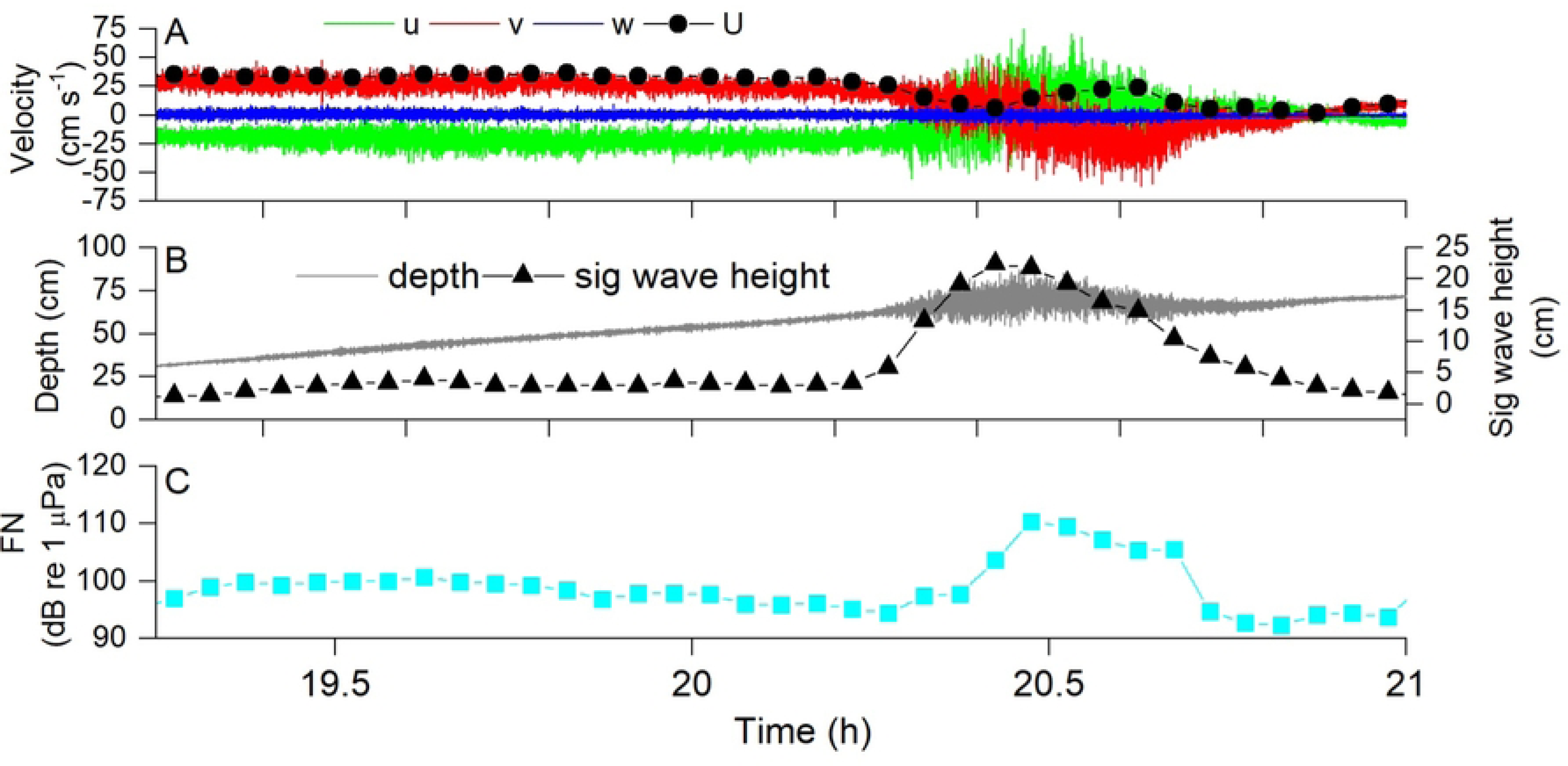
Example data from the mudflat showing the impact of a severe thunderstorm (occurring approximately at time 20.25 - 20.75) on (A) flow velocity, (B) significant wave height H_S_, and (C) FN. Note higher FN at the beginning of the tide when velocity was high, then a spike in FN coinciding with a massive increase in H_s_ with the thunderstorm, followed by lower FN after the storm when velocity was lower. This example demonstrates how we were able to correlate time-averaged values of hydrodynamic parameters to the soundscape. In this example 3 min averaging was used to better demonstrate the impact of the storm, but all other results utilized 15 min averaging in order to allow sufficient time to calculate turbulence statistics. There were high amounts of rain during this period, which can also be detected at these frequencies (25).

Flow and wave characteristics were similar at the three sites. Maximum flow speeds were 21 cm s^-1^ at the natural reef, 32 cm s^-1^ at the restored reef, and 37.6 cm s^-1^ at the mudflat. Significant wave height (H_s_) was relatively low at the two reefs, with 15 min averages ranging from 0.1 - 4.4 cm at the natural reef and 0.2 - 3.8 cm at the restored reef. Mudflat measurements were conducted during windier conditions, and as a result H_s_ was much greater, ranging from 0.5 - 18 cm.

ADV data allows for the calculation of several turbulence statistics, which in well-developed boundary layer flow should all be highly correlated to one another (Reidenbach et al. 2006). Indeed, Fig. 5 shows that turbulent kinetic energy (TKE) and turbulence dissipation (ε) were closely related, as were U and TKE. Both the slopes and strengths of these correlations were substantially greater at the reef sites than at the mudflat due to larger benthic roughness at the reefs (6).

**Figure 5.**
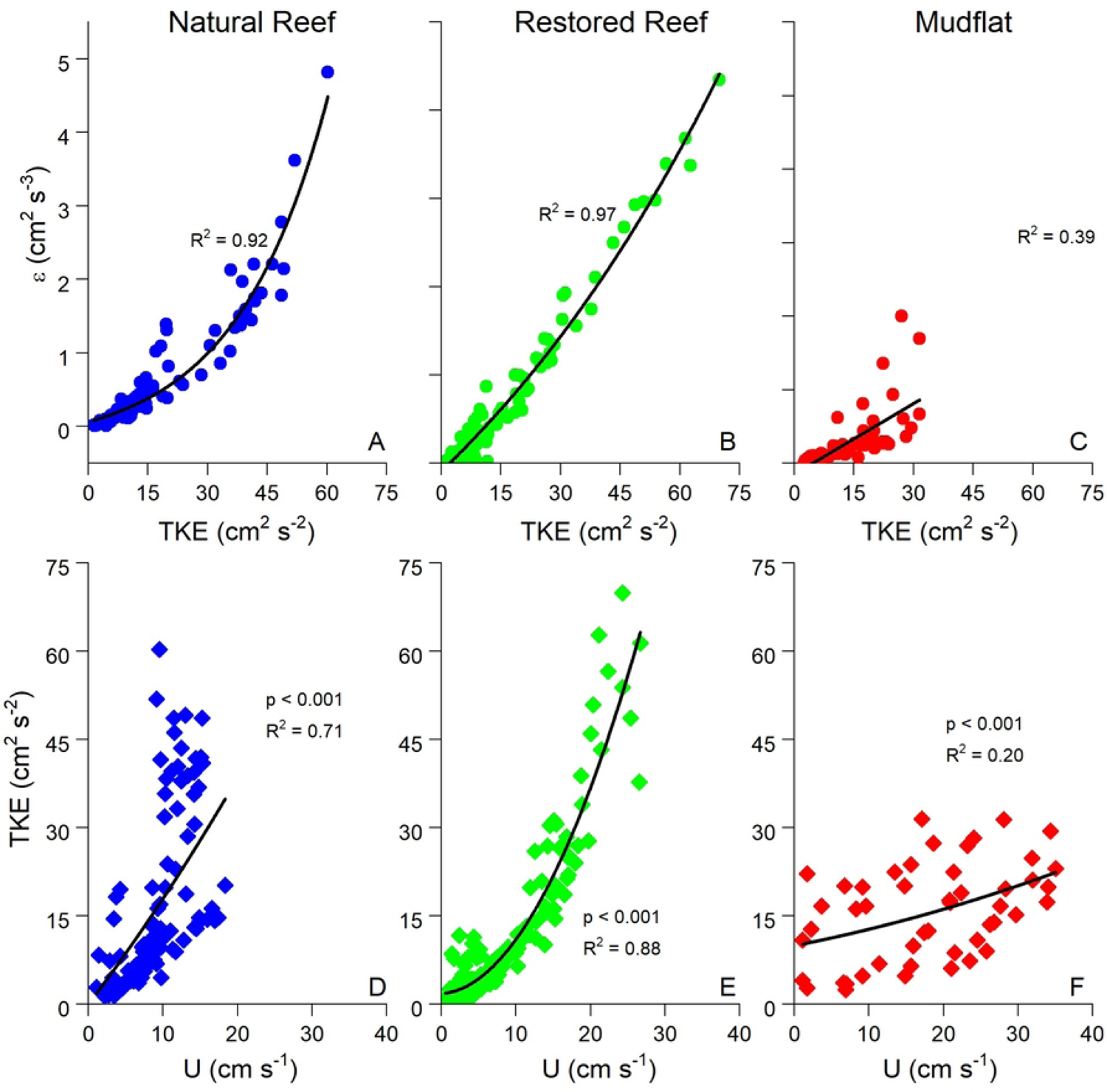
(A, B, C) Exponential fits of turbulent kinetic energy (TKE) vs. turbulence dissipation (ε) and (D, E, F) linear fits of flow speed (U) vs. TKE. TKE vs. ε was similar at all three sites, but the U vs. TKE slope was significantly steeper at the reef sites than the mudflat due to higher benthic roughness. Each datapoint represents a 15 min average. Only data with significant wave height < 5 cm were used at the mudflat site in order to avoid possible wave artifacts.

### 4.3 Turbulence and flow noise

Fig. 6 compares FN with U, TKE, and ε across all three sites, with all exhibiting exponential relationships. These relationships were relatively weak when all data were included (R^2^ = 0.11 – 0.26) indicative of the range of flow and turbulence conditions at the three sites that varied due to ebbing and flooding conditions. The statistical relationship between FN and flow parameters became much stronger when looking at individual submerged periods at a single site as shown in Fig. 7. Here, exponential fits of FN vs. U, TKE, and ε over a representative submerged tidal period at the natural reef had R^2^ = 0.63 - 0.98. FN was also weakly correlated with significant wave height at all three sites, indicating the detection of non-breaking wave orbitals within the soundscape and the generation of turbulence due to oscillatory wave motion (Fig. 8).

**Figure 6.**
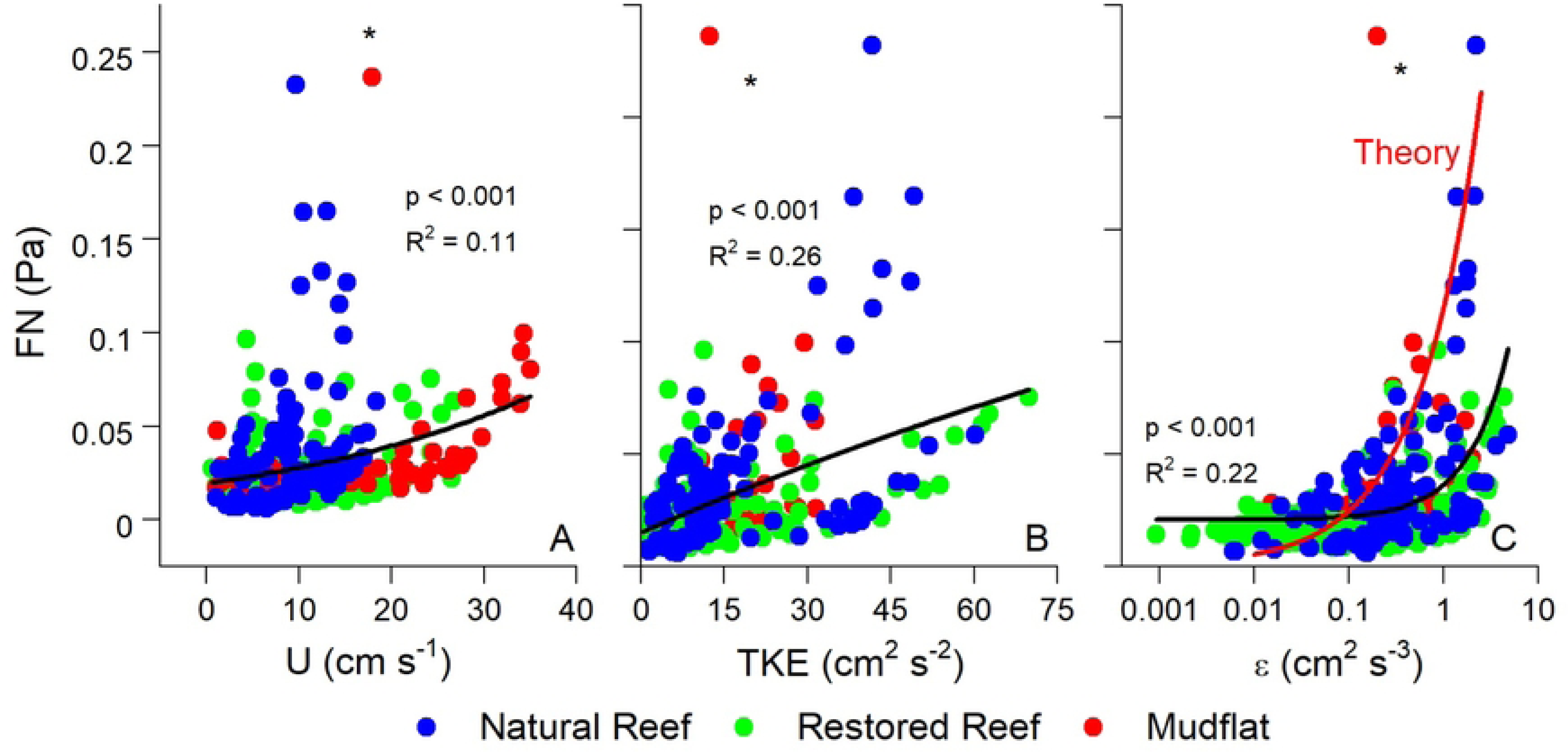
(A) Flow speed (U), (B) turbulent kinetic energy (TKE), and (C) turbulence dissipation (ε) vs. FN at the three sites. These data were fit with exponential functions. Panel C also shows the theoretical FN vs. ε relationship predicted by Eq. 10-12. Both TKE and ε had stronger effects on FN than U, as water column turbulence results in pressure fluctuations that are recorded by the hydrophone. An outlier point from the mudflat (asterisk) was not included. Please note for this comparison FN has been converted from dB to Pa.

**Figure 7.**
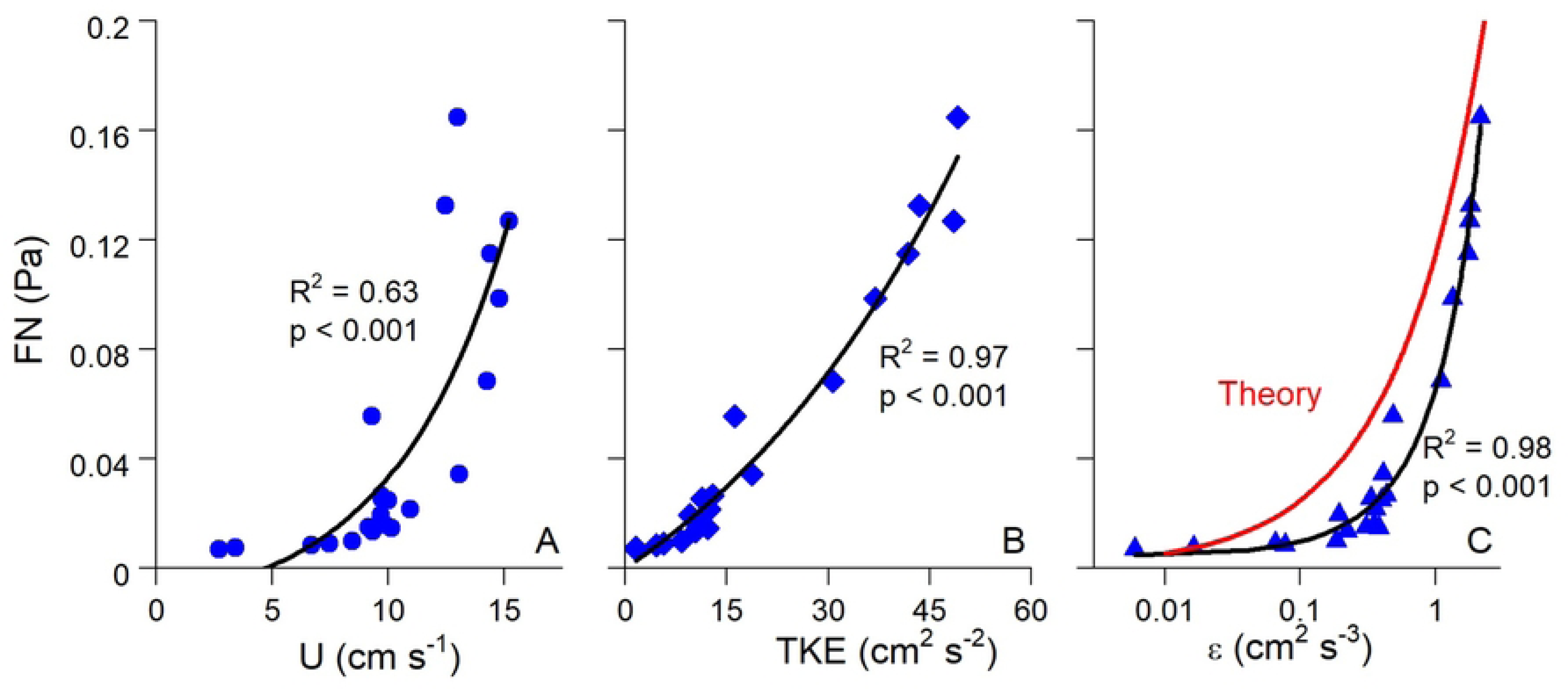
(A) Flow speed (U), (B) turbulent kinetic energy (TKE), and (C) turbulence dissipation (ε) vs. FN over a representative submerged high tide period at the natural reef. Panel C also shows the theoretical FN vs. ε relationship predicted by Eq. 10-12. Although correlations between FN and hydrodynamic parameters were relatively weak when integrating over all data (Fig. 6), this effect is likely due to differences in background noise between different submerged periods. During each submerged period, which better controls for background noise levels, correlations between FN and hydrodynamics were much stronger. Please note for this comparison FN has been converted from dB to Pa.

**Figure 8.**
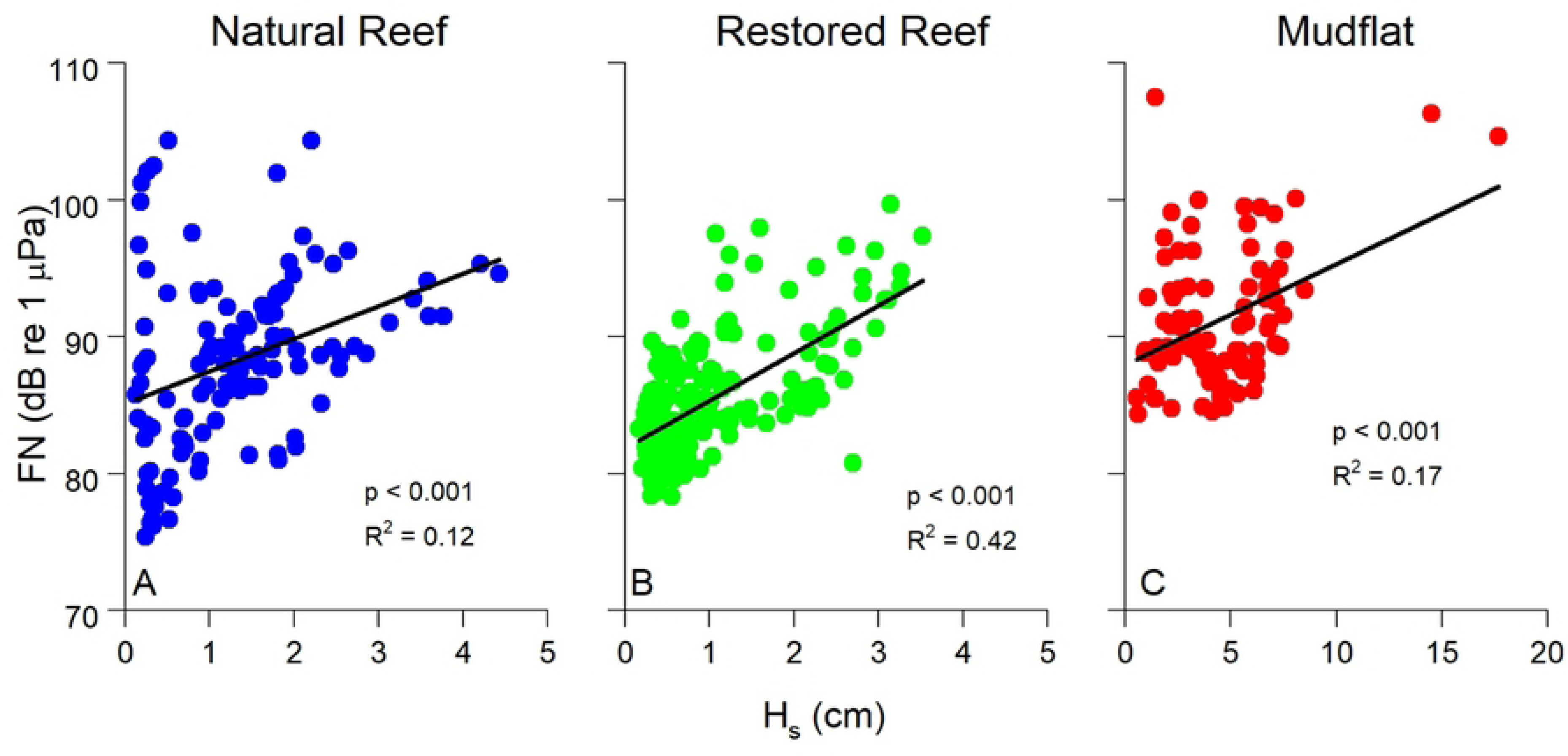
Significant wave height (H_s_) vs. FN at all three sites. H_s_ was significantly (p < 0.001) related to FN at all three sites, showing the impact of wave orbitals on low frequency pressure fluctuations recorded by the hydrophone. Note the different scale at the mudflat (C), as conditions were significantly wavier during measurements at this site. The two datapoints at high values of H_s_ in (C) represent the storm depicted in Fig. 3. Data were averaged over 15 min intervals.

Fig. 6C and 7C depict the theoretical FN vs. ε relationship predicted by Eq. 9-11. We set the characteristic eddy size to 0.1 m in this calculation to account for the height of the ADV above the reef, as explained above. Despite the slightly different curve shapes, Fig. 6C and 7C show close quantitative agreement between theory and measurements. This suggests that the low frequency noises that we measured are correctly designated as FN, arising in large part by the local turbulence in the system. Interestingly, the theoretical relationship did a better job at explaining the highest FN values than our exponential best-fits, as shown in Fig. 6C.

We computed the bed shear stress *τ_b_* from ε for each site using eq. 4, with FN and τ_b_ of similar magnitude (Fig. 9). These results suggest that shear stresses derived from flow interaction with the seafloor scales with pressure fluctuations within the FN (f < 100 Hz) sound spectrum, and that the dominant mechanism in producing near-field sound is bed-generated turbulence.

**Figure 9.**
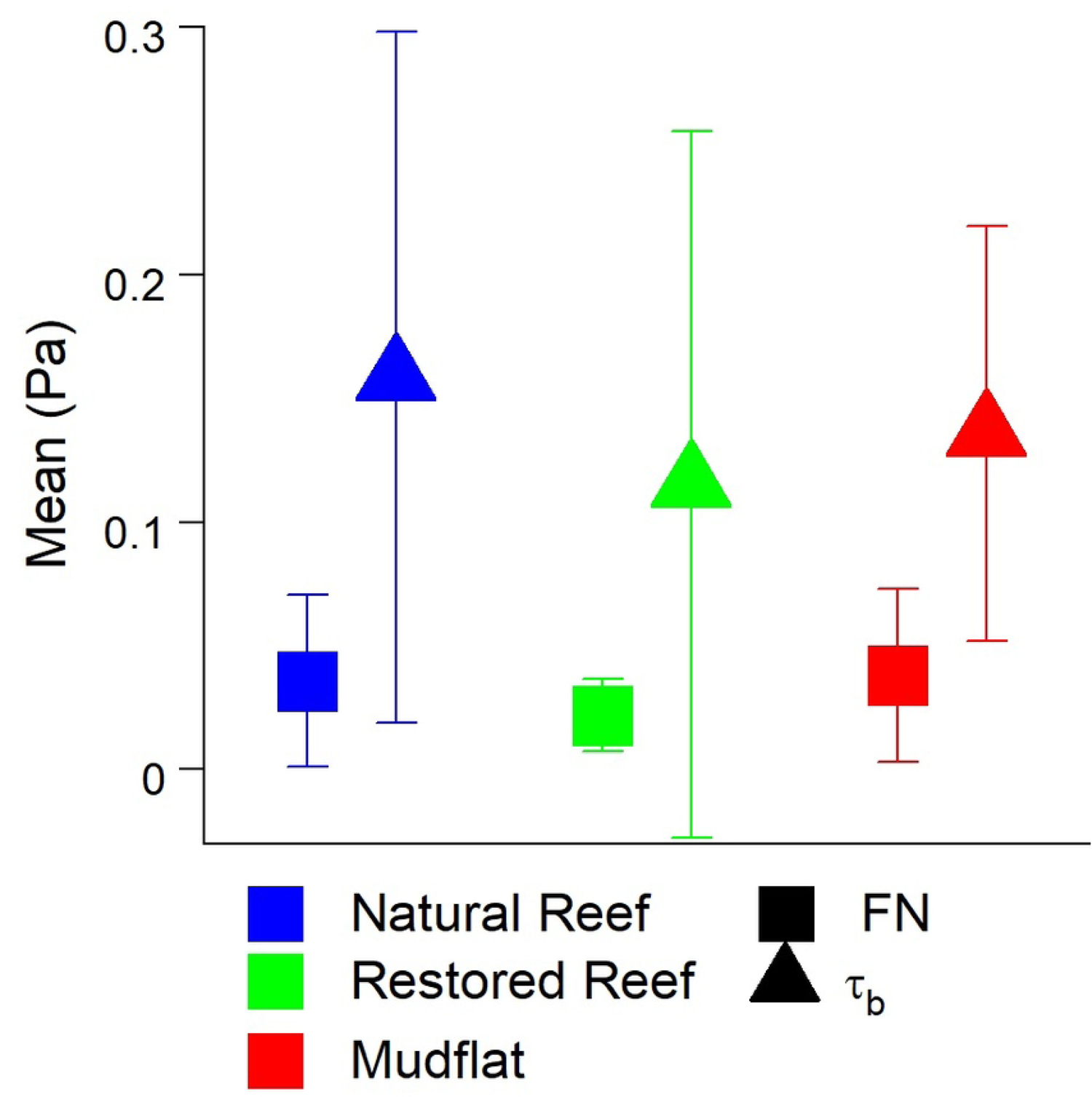
Mean (A) FN and (B) bed shear stress (τ_b_) at all three sites ± SD. If FN is caused in large part by turbulent pressure fluctuations, it should be of similar magnitude as τ_b_ (Vollmer & Kleinhans 2007). Please note for this comparison FN has been converted from dB to Pa.

## 5 Discussion

The governing equations of fluid dynamics are the Navier-Stokes (N-S) equations. For three-dimensional incompressible flow, the equation for x-momentum yields:

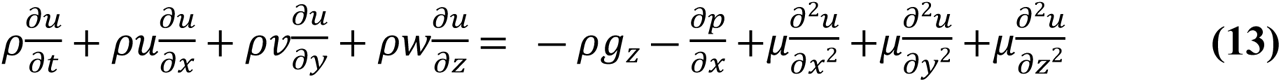

where x, y, and z are the horizontal, transverse, and vertical directions, respectively, ρ is density, *g*_z_ is the force of gravity in the z direction, p is pressure, μ is dynamic viscosity, and u, v, w are the velocities in the x, y, and z directions. According to N-S, changes in the velocity field are directly related to changes or fluctuations in pressure, and as a result turbulent motions in the water column generate fluctuations in pressure. These fluctuations are detectable by pressure-sensing instruments such as hydrophones, a fact first demonstrated by Wenz (1962) from the analysis of the spectral signature of hydrophone recordings. In more recent years this “flow-noise” (FN) or “pseudosound” has been more frequently studied, often with the goal of removing these effects from the soundscape as they do not represent true sound waves ( Bassett et al. 2014; von Benda-Beckmann et al. 2016). However, as we show in this study FN contains a wealth of information about the physical environment, and should not necessarily be discarded from soundscape studies.

Correlations between FN and hydrodynamics were much stronger over individual submerged periods (R^2^ = 0.63 – 0.98; Fig. 7) than for combined data (R^2^ = 0.11 - 0.26; Fig. 6), a pattern that may be attributed to differences in the background acoustic environment across different submerged high tide periods, perhaps due to wind or rain (Ma et al., 2005). Additionally, correlations between TKE and ε with FN were stronger than those between U and FN. These results show that turbulence is a better predictor of FN than mean flow speed, agreeing with past acoustic measurements conducted in air flow (50). Turbulent motions also generate pressure fluctuations near the bed (51). For aquatic systems this predicted relationship is *p*_*rms*_ = 3.1 *τ*_*b*_ (52), where *p*_*rms*_is the root mean square of these fluctuations and is equivalent to FN. In our study τ_b_ was invariably greater than FN, likely due to the fact that not all scales of pressure fluctuations were measured by the hydrophone. Nevertheless, FN fell within the range of ± 1 SD of *τ*_*b*_(Fig. 9), demonstrating reasonable agreement.

Best fit relationships for FN vs. ε closely agreed with the theoretical fit predicted by Eq. 9-11 (Fig. 6C, 7C), with theory best at explaining the highest values of FN (Fig. 6C). FN in our study was defined as the mean of the pressure fluctuations recorded by the hydrophone at *f* < 100 Hz. Included in this range, *f* < 100, are non-breaking wave orbitals (Fig. 9), ultra-low frequency noises such as ships (24), and the contribution of self-noise of the hydrophone, all of which are difficult to separate from ambient turbulence. Additionally, despite having the lowest levels of turbulence, the mudflat had the highest mean FN among the three sites indicating that factors other than turbulence can impact our results. Nevertheless, the strong agreement we observed between predicted and actual FN vs. ε relationships is very promising, indicating that the bulk of FN can be attributed to turbulent pressure fluctuations although future studies should be conducted to better elucidate this relationship. Both burying the hydrophones to diminish the impact of self-noise, as well as using hydrophones tuned to capture ultra-low frequencies would likely result in FN vs. ε results that more closely match our theory.

In terms of reef sounds, ranges for SPL_MID_ and SPL_HI_ (∼90-100 dB; Fig. 3) both fell within the range of past values measured over subtidal oyster reefs (18). While past studies have shown relatively similar SPL_MID_ between reefs and unstructured bottoms, there is expected to be a wide range of variability between SPL at frequencies > 2 kHz (18,37). SPL_HI_ was in fact the greatest at the restored reef, but approximately equal at the natural reef and mudflat. This may be due in part to the relatively small size of the natural reef (240 m^2^), which was approximately 1/10^th^ the size of the restored reef (2275 m^2^), resulting in less reef area, and potentially fewer organisms, that contributed to the recorded sound. This spatial disparity has important implications for larval recruitment, as oyster patch size affects both water column turbulence and sounds detected by pelagic oyster larvae (37,53), with larger patches predicted to enhance settlement success.

Pelagic oyster larvae use water column turbulence as a settling cue, as increases in local acceleration resulting from turbulence induce downward swimming behavior (Fuchs et al., 2013, 2015; Wheeler et al., 2015). Fuchs et al. (2013) showed that ε was the dominant turbulent cue for downward larval swimming velocity (w_b_). In our study, ε was also a dominant hydrodynamic indicator of FN (Fig. 6, 7), a fact that has been previously demonstrated in tidal channels (29). Laboratory experiments of swimming behavior of competent oyster larvae found larvae to actively swim downward in response to turbulence dissipation rates ε > 0.1 cm^2^ s^-3^ (Fuchs et al. 2013). Coincidentally, ε = 0.1 cm^2^ s^-3^ is approximately where our observed and theoretical FN vs. ε are equal, with FN ≈ 0.025 Pa. Both empirical and predicted relationships predict rapid increases in FN above ε = 0.1 cm^2^ s^-3^, with theory predicting the sharper increase (Fig. 6C), suggesting that larvae alter swimming behaviors in response to FN ≥ 0.025 Pa within our oyster reef system. Fuchs et al. (2013) reported a similar relationship between ε and w_b_ as we observed between ε and FN, showing an inflection point with substantial increases in w_b_ when ε > 0.1 cm^2^ s^-3^. Taken together with our study, these results imply that downward swimming behavior of settling larvae may be proportional to FN once a minimum ε of 0.1 cm^2^ s^-3^ (or FN = 0.025 Pa) is reached.

The organ used by many molluscan species to detect sound is the statocyst (34), which consists of a mineralized mass within a ciliated sac (55). Not only do oyster larvae use these organs to detect turbulence (Fuchs et al., 2015), but adult oysters likely use them to respond to low frequency sounds in the range of turbulent pressure fluctuations (10-80 Hz; Charifi et al., 2017). It is therefore possible that larval settlement induced by sound and turbulence do not result from differing responses to separate stimuli, but rather are both equivalent statocyst-driven responses to turbulent pressure fluctuations “heard” by the larvae. Past studies of oyster reef soundscapes have inferred that higher frequency (i.e. *f* > 2000 Hz) sound induces larval settlement by describing differences between reefs and unstructured bottom at these frequencies (36,37), however these studies did not examine lower frequencies (*f* < 100 Hz) in the range of FN. Likewise, neither Fuchs et al. (2013) nor Wheeler et al. (2015) examined the impact of turbulent pressure fluctuations on larval settlement, only velocity effects. In fact, as stated above Fuchs et al. (2013) found that ε was the only turbulent cue that larvae consistently responded to, even in low-turbulence conditions.

Besides its impact on larval settlement, turbulence is an important control of many oyster reef processes, including whole-reef metabolism, dissolved and particulate fluxes, sediment transport (Reidenbach et al. 2013, Volaric et al. 2018), and oyster growth and mortality (Lenihan, 1999). Detailed measurements of turbulence are expensive, require a high degree of technical expertise, and are often difficult to conduct over longer timeframes (i.e. weeks to years). In recent years, bioacoustic monitoring of marine ecosystems has shown promise in describing their overall health and biodiversity (12,18). Further research into linking the soundscape to hydrodynamics in these endeavors would help to better describe how physical controls impact biological activity, further aiding future monitoring efforts of aquatic ecosystems.

## 6 Conclusions

In this study we made side-by-side hydrophone and ADV measurements over two intertidal oyster reefs and one intertidal mudflat, finding that FN was correlated to important hydrodynamic parameters such as U, TKE, ε, *τ*_*b*_, and H_s_. SPL varied significantly across the sites, with highest SPL_MID_ and SPL_HI_ for the restored reef. Our results show a similar relationship between ε and FN that has previously been shown for ε and larval settling velocities. Reef sounds also attract these pelagic larvae, a similarity that might be explained by the statocyst. This pressure-sensitive organ is biologically analogous to a hydrophone and is used by mollusks such as oysters to detect both sound and turbulence. Our results have important implications for understanding larval recruitment and demonstrate possible applications for ultra-low frequency (*f* < 100 Hz) data often discarded in soundscape studies.

## Acknowledgements

We would like to thank B. Lusk and The Nature Conservancy for providing access to the sites, A. Turgut and O. Caretti for research guidance, and D. Lee and C. Johnston for field assistance. Support for this study was provided by the National Science Foundation (NSF) through grants for the Virginia Coast Reserve Long-Term Ecological Research program (DEB-1237733 and DEB-1832221). Support was also provided by the Coastal Futures Conservatory (http://www.coastalconservatory.org/), and by the University of Virginia. SA was funded in part by NCI U01 CA227544 awarded to Herbert Sauro and H. Steven Wiley. The authors declare no conflicts of interest.

## Data Availability Statement

The data presented in this study are openly available via the Environmental Data Initiative data portal at https://doi.org/10.6073/pasta/3a372383f1bbaf8367232adc592e135e.

